# Oleic acid-related anti-inflammatory effects in force-stressed PdL fibroblasts are mediated by H3 lysine acetylation associated with altered *IL10* expression

**DOI:** 10.1101/2022.01.11.475909

**Authors:** Lisa Schuldt, Katrin von Brandenstein, Collin Jacobs, Judit Symmank

## Abstract

The initiation of a spatially and temporally limited inflammation is essential for tissue and bone remodeling by the periodontal ligament (PdL) located between teeth and alveolar bone. Obesity-associated hyperlipidemic changes may impair PdL fibroblast (PdLF) functions, disturbing their inflammatory response to mechanical stress such as those occurring during orthodontic tooth movement (OTM). Recently, we reported an attenuated pro-inflammatory response of human PdLF (HPdLF) to compressive forces when stimulated with monounsaturated oleic acid (OA). Fatty acids, including OA, could serve as alternative source of acetyl-CoA, thereby affecting epigenetic histone marks such as histone 3 lysine acetylation (H3Kac) in a lipid metabolism-dependent manner. In this study, we therefore aimed to investigate the extent to which OA exerts its anti -inflammatory effect via changes in H3Kac. Six-hour compressed HPdLF showed increased H3Kac when cultured with OA. Inhibition of histone deacetylases resulted in a comparable *IL10* increase as observed in compressed OA cultures. In contrast, inhibition of histone acetyltransferases, particularly p300/CBP, in compressed HPdLF exposed to OA led to an inflammatory response comparable to compressed control cells. OA-dependent increased association of H3Kac to *IL10* promoter regions in force-stressed HPdLF further strengthened the assumption that OA exhibits its anti-inflammatory properties via modulation of this epigenetic mark. In conclusion, our study strongly suggests that obesity-related hyperlipidemia affect the functions of PdL cells via alterations in their epigenetic code. Since epigenetic inhibitors are already widely used clinically, they may hold promise for novel approaches to limit obesity-related risks during OTM.

## Introduction

Non-communicable diseases (NCDs) such as cancer, cardiovascular disease or diabetes mellitus are the main cause of premature mortality [1]. As a chronic progressive disease, obesity substantially increases the risk of some cancers, heart attacks and strokes, fatty liver disease, type 2 diabetes mellitus, and osteoarthritis, as well as mental health conditions such as dementia and depression, thereby promoting the progression of NCDs [1]. However, the causes of obesity are exceedingly complex and involve, in addition to an obesogenic environment, a genetic and epigenetic background [2]. In this regard, various studies have demonstrated that under obese conditions, acetylation levels of certain histone lysines are altered [3–6]. This dynamic epigenetic modification alters chromatin architecture and regulates the accessibility of genes [7]. Histone acetyltransferases (HAT), known as epigenetic writers, perform acetylation of specific lysines of histone tail proteins, for which acetyl-CoA is required [8]. Epigenetic erasers in this process are histone deacetylases (HDACs), which specifically remove acetylation marks on histone lysines [7].

Numerous studies on the underlying mechanisms of obesity-related systemic and organ-specific changes have focused on the effects of increases in serum free fatty acids (FFA) [9–14]. Under healthy conditions, FFA are important for normal cellular functions and could serve as alternative source of acetyl-CoA in histone acetylation processes [15,16]. However, hyperlipidemia can impair these processes and thus alter cell-specific gene expression [17–24]. While saturated fatty acids (SFA) have been shown to stimulate the expression of genes encoding pro-inflammatory cytokines such as tumor necrosis factor α (TNFα), interleukin 6 (IL6), IL8, IL1α, and IL1β [17–22,25,26], monounsaturated fatty acids (MUFA) are mostly considered to have anti-inflammatory effects [11,20,23,24,27]. However, both types of FFA were found increased in serum of obese patients [12,15,28]. Among those MUFA, mono-unsaturated oleic acid (omega-9 fatty acid; 18:1; OA), which is also highly enriched in the Mediterranean diet, has been shown to counterbalancing the pro-inflammatory effects of SFA [11,20,24,27]. In this context, OA-related increased anti-inflammatory IL10 secretion was reported [29], contributing to the down-regulation of pro-inflammatory cytokines [30,31]. Moreover, there is evidence that an OA-enriched diet contributes to the treatment and prevention of obesity, as it can lead to a reduction in abdominal fat and thus positively influence body composition [32].

We recently reported that OA in contrast to the SFA palmitic acid (PA) attenuates the force-induced pro-inflammatory response of human periodontal ligament fibroblasts (HPdLF) [33], which are the main cell type of the periodontal ligament (PdL). As connective tissue between the teeth and the alveolar bone, the PdL has important functions in modulating the response of soft and hard tissues to mechanical stimuli, such as those that occur during trauma and mastication, but also in orthodontic treatments [34]. During tissue remodeling caused by mechanical stimulation, HPdLF promote transient aseptic inflammation, which is essential for tissue and bone remodeling processes [35]. Interfering with inflammatory signaling, both by decreasing and increasing the inflammatory response of HPdLF may affect tissue and bone remodeling increasing the risks for attachment loss, root resorption and tooth loss [36,37].

The impact of increased body mass index (BMI) on orthodontic tooth movement (OTM) in children and adolescents is, however, controversial showing both reduced and increased OTM rates in different studies [38–41]. Obesity exhibits a very heterogeneous clinical pattern, which could explain the different findings, since only the BMI was used for obese classification. To date, there are few *in vitro* studies addressing the underlying biological mechanisms in regard to obesity-associated hyperlipidemic conditions. However, the reasons explaining why OA exerts anti-inflammatory effects on the force-induced response of HPdLF that may influence bone remodeling processes remained unclear. In view of the potential effect of increased OA exposure on acetyl-CoA levels and thus on histone acetylation, our aim was to analyze to what extent this contributes to its anti-inflammatory effect in compressed HPdLF. Thus, the results of our study should contribute to the understanding of how metabolic changes due to hyperlipidemic states affect the epigenetic regulation of inflammatory markers in the PdL and thus may have implications for the force-induced response of PdL fibroblasts.

## Materials and Methods

### Cell culture

Commercially acquired human periodontal ligament fibroblasts (HPdLF, Lonza, Basel, Switzerland) are pooled from several donors. They were grown in Dulbecco’s modified Eagle medium (DMEM; Thermo Fisher Scientific, Carlsbad, CA, USA) containing 4.5 g/L glucose, 10 % heat-inactivated fetal bovine serum (Thermo Fisher Scientific, Carlsbad, CA, USA), 100 U/ml penicillin, 100 μg/ml streptomycin and 50 mg/L L-ascorbic acid at 37 °C, 5% CO2 and 95% humidity. When reaching 75% confluency, HPdLF were passaged with 0.05% Trypsin/EDTA (Thermo Fisher Scientific, Carlsbad, CA, USA). Passage four to eight were used for experimental setups. For transcriptional analysis, HDAC and HAT activity assay as well as CHIP, 2.5 × 10^4^ HPdLF were seeded into each well of a 6-well plate. For TPH1 adhesion assay, 5 × 10^3^ HPdLF were seeded on glass coverslips into each well of a 24-well-plate.

THP1 cells (DMSZ, Braunschweig, Germany) were cultured in RPMI 1640 medium (Thermo Fisher Scientific, Carlsbad, CA, USA) that contains 10% FBS, 100 U/ml penicillin and 100 μg/ml streptomycin at 37 °C, 5% CO2 and 95% humidity. Passages were performed weekly and 1 x 10^6^ cells were reseeded into a T175 culture flask (Thermo Fisher Scientific, Carlsbad, CA, USA).

### Fatty acid exposure

Prior treatment with 200 μM oleic acid (OA) for six days, freshly seeded HPdLF were subsequently cultured for 24h in culture medium. The stimulation with OA was performed as described previously [42]. Briefly, OA was dissolved at 70 °C in sterile water containing 50 mM NaOH, complexed with 37 °C preheated bovine serum albumin (BSA, Seqens IVD, Limoges, France) and diluted in culture medium. BSA (0,66%) was used as control.

### Inhibitor treatment

Different concentrations of romidepsin between 10 nM and 100 nM were used to inhibit histone deacetylases in 75% confluent HPdLF. For inhibition of histone acetyltransferases, 1 nM to 10 nM anacardic acid (AA) or 1 nM to 10 nM C646 were applied. For final experiments, 10 nM romidepsin, 10 nM AA or 1 nM C646 were used. Since the inhibitors were dissolved in DMSO, this was used as control.

### Mechanical compression

HPdLF were stimulated by compressive force as previously described [33]. Briefly, sterile glass plates weighing 2 g/cm^2^ were placed in 6-well plates for six hours at 37 °C, 5% CO2, and 95% humidity. In 24-well plates, the application of compressive force was performed by centrifugation for six hours at 37 °C and a force of 7.13 g/cm2, which was the minimal condition of the centrifuge.

### THP1 cell adherence assay

For visualization of the inflammatory response of specifically stimulated HPdLF, analysis of THP1 cell adhesion was performed as previously described [33]. Briefly, after staining of non-adherent cells with 15 μM Celltracker CMFDA (Thermo Fisher Scientific, Carlsbad, CA, USA) for 30 min at 37 °C, 50 × 103 cells were added to each well of cultured HPdLF in a 24-well plate. After cell adhesion for 30 min at 37 °C, 5% CO2, and 95% humidity, non-adhered THP1 cells were removed with prewarmed PBS and remaining cells were fixated in 4% paraformaldehyde for 10 min. Cell nuclei were stained for 5 min with DAPI (1:10,000 in PBS) and coverslips were embedded with Mowiol^®^ 4–88 (Carl Roth, Karlsruhe, Germany) on glass object slides for microscopic imaging.

### RNA extraction and quantitative PCR

TRIzol Reagent (Thermo Fisher Scientific, Carlsbad, CA, USA)/1-bromo-3-chloropropane was used to isolate RNA of specifically treated HPdLF for expression analysis. After purification with the RNA Clean and Concentrator-5 kit (Zymo Research, Freiburg, Germany), quantity and quality was tested with Nanodrop 2000 (Avantor, Radnor, PA, USA). cDNA synthesis was performed with SuperScript IV Reverse Transcriptase (Thermo Fisher Scientific, Carlsbad, CA, USA) and Oligo(dt)18 primers (Thermo Fisher Scientific, Carlsbad, CA, USA), according to the manufacture’s protocol. Luminaris Color HiGreen qPCR Master Mix (Thermo Fisher Scientific, Carlsbad, CA, USA) and the qTOWER3 (Analytik Jena, Jena, Germany) was used for performing quantitative PCR. All analyzed genes are displayed in **Table 1** with their respective primer sequences. *RPL22* and *TBP* were used as reference genes. To evaluate primer quality and specificity, melting curve analysis and agarose gel electrophoresis were performed. Primer efficiency was analyzed with a cDNA dilution series. The efficiency corrected ΔΔCT method was used for data analysis [43]. Each condition was analyzed at least in biological triplicates with technical duplicates.

**Table 1.**
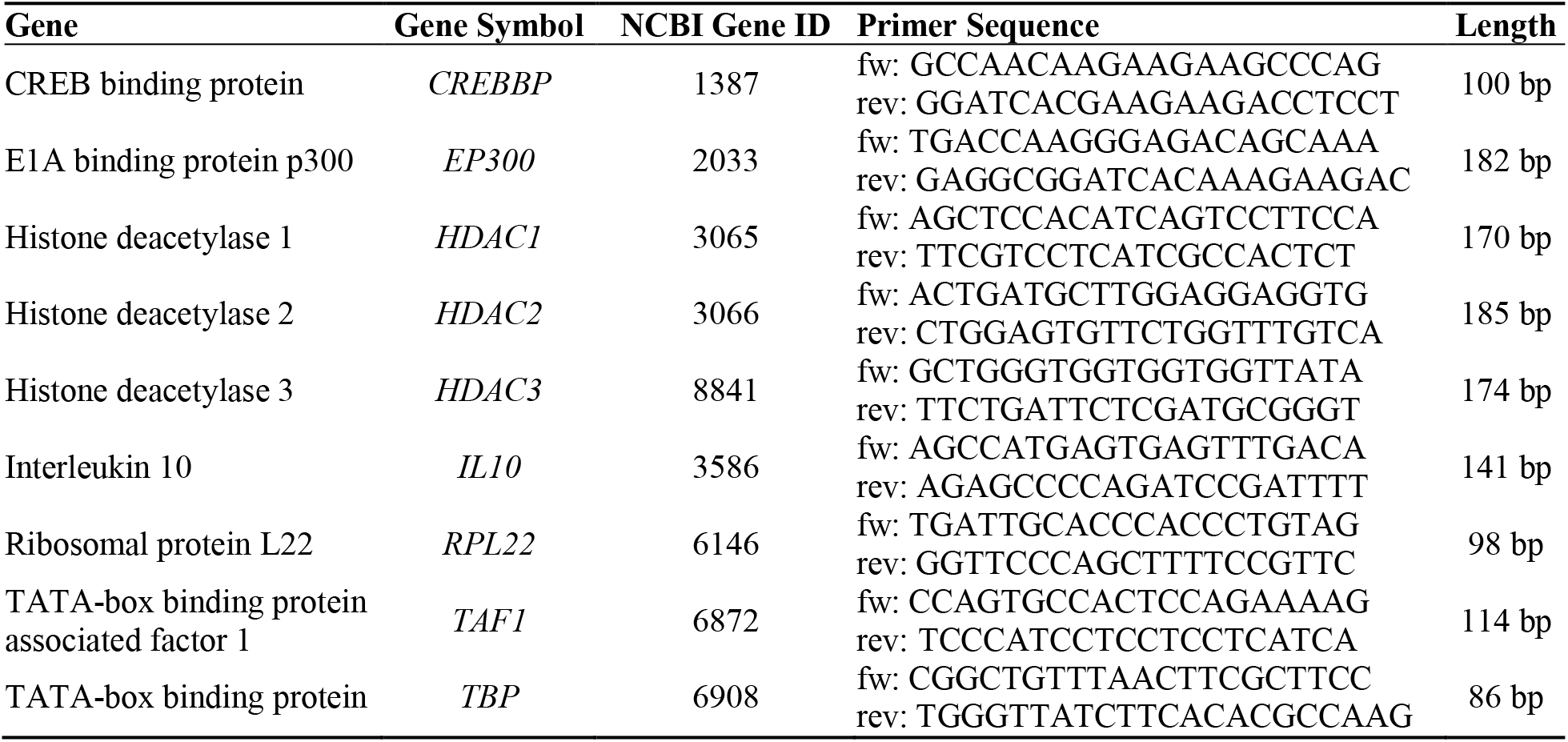
qPCR primer sequences of human genes indicated in 5’ −3’ direction. bp, base pairs; fw, forward; Length, amplicon length; rev, reverse.

### Immunofluorescent staining

After specific treatment, HPdLF cultured on coverslips were fixed in 4% PFA for 10 minutes and washed with PBS before incubation with the primary and secondary antibodies for 45 minutes each. Nuclei were stained with DAPI (Thermo Fisher Scientific, Carlsbad, CA, USA; 1:10000 in PBS). Mouse-anti-human H3K9/14/18/23/27ac (Thermo Fisher Scientific, Carlsbad, CA, USA; 1:500) was used to detect this histone modification in HPdLF and goat-anti-Mouse-Cy5 (Jackson ImmunoResearch, XXX; 1:1000) was used for fluorescently labeling. Each condition was tested in independent biological triplicates with technical duplicates.

### MTT Cell Vitality Test

Cell vitality of HPdLF was analyzed with MTT colorimetric assay (Sigma Aldrich, St. Louis, Missouri, USA) according to manufacturer’s protocol. Colorimetric analysis was performed with the plate reader Infinite ^®^ M Nano (TECAN, Männedorf, Swiss). Each condition was analyzed in biological triplicates with technical duplicates.

### Nuclear Extraction and HDAC/HAT Activity Assay

Nuclear extraction was performed with the EpiQuik Nuclear Extraction Kit (EpiGentek, Farmingdale, New York, USA) according to manufacturer’s protocol. The activities of histone deacetylases (HDAC) and histone acetyltransferases (HAT) were analyzed with EpiQuik HDAC Activity/Inhibition Assay Kit (EpiGentek, Farmingdale, New York, USA) and EpiQuik HAT Activity/Inhibition Assay Kit (EpiGentek, Farmingdale, New York, USA) according to manufacturer’s guidelines, respectively. Assay read out was done with the plate reader Infinite ^®^ M Nano (TECAN, Männedorf, Swiss). Each condition was analyzed in biological triplicates with technical duplicates.

### Chromatin Immunoprecipitation (ChIP)-qPCR

After treatment, crosslinking of DNA and protein was performed with 1% formaldehyde in PBS for 10 min and neutralized with 120 mM glycine. Cells were harvested in ice-cold PBS, aliquoted with an approximate cell number of 1 × 10^6^ cells per tube and centrifuged for 5 min at 1000 x g at 4°C prior storage at −80 °C. Chromatin immunoprecipitation was performed with the Zymo-Spin ChIP Kit (Zymo Research, Freiburg, Germany) according to manufacturer’s protocol Following ChIP-valuated antibodies were used for precipitation: mouse-anti-human H3K9/14/18/23/27ac (Thermo Fisher Scientific, Carlsbad, CA, USA) and mouse-anti-IgG (Thermo Fisher Scientific, Carlsbad, CA, USA). One percent input controls were stored throughout the ChIP process. The determination of the amount of specific DNA fragments bound to H3K9/14/18/23/27ac was performed as previously described [44]. Briefly, a 20-cycle primer-specific pre-amplification was performed prior quantitative analysis with the qTOWER3 (Analytik Jena, Jena, Germany) according to the manufacturer’s protocol using Luminaris Color HiGreen qPCR Master Mix (Thermo Fisher Scientific, Carlsbad, CA, USA). Primer sequences for three promoter regions (#1, #2 and #3) as well as one non-promoter region (#4) are displayed in **Table 2**. Normalization of DNA content was performed according to the per cent-input method in relation to the analyzed input probes [45]. Each condition was analyzed in biological triplicates with technical duplicates.

**Table 2.**
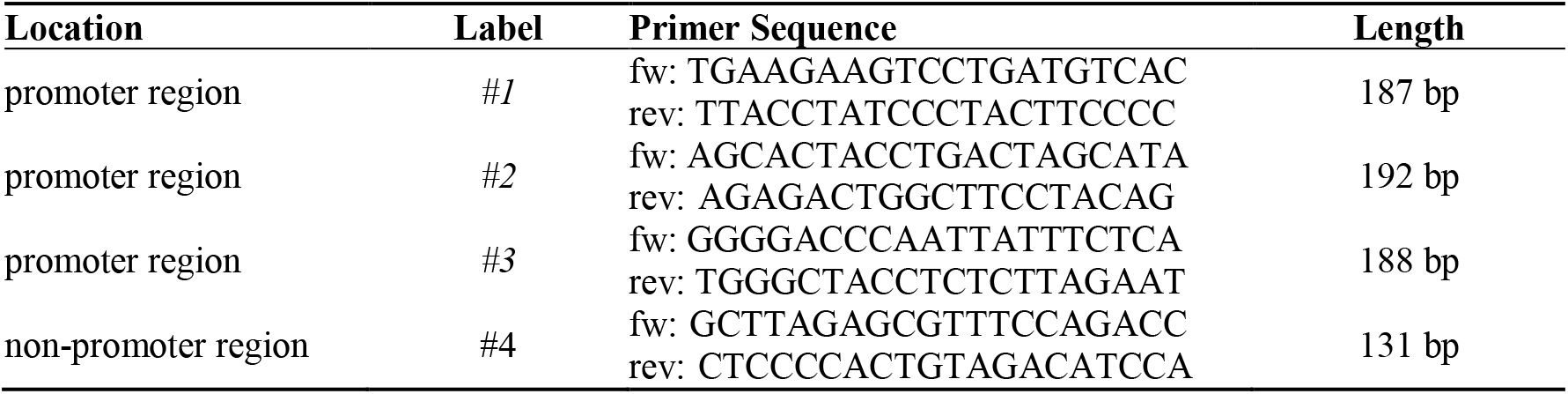
qPCR primer sequences of human *IL10* in promoter regions (#1, #2, #3) and a non-promoter region (#4) indicated in 5’ −3’ direction. bp, base pairs; fw, forward; Length, amplicon length; rev, reverse.

### Microscopy, Image Analysis and Statistics

The THP1 adhesion assay and the H3K9/14/18/23/27ac immunofluorescent staining was imaged with the inverted confocal laser scanning microscope TCS SP5 (Leica, Wetzlar, Germany). Analysis of the THP1 cell number and the fluorescent intensity of H3K9/14/18/23/27ac staining was performed with Fiji software (https://imagej.net/Fiji, accessed on 01.04.2017). For statistical analysis, Graph Pad Prism (https://www.graphpad.com, accessed on 01.02.2021). Adobe Photoshop CS5 (https://adobe.com, accessed on 01.02.2013) was used for figure illustration. One-way ANOVA and post hoc test (Tukey) were used as statistical tests. Significance levels: p value < 0.05 */#/§; p value < 0.01 **/##/§§; p value < 0.001 ***/###/§§§

## Results

### Oleic acid exposure enhanced the force-related up-regulation of H3 lysine acetylation

To determine the influence of oleic acid (OA) on force-induced changes in H3 lysine methylation, we first examined the expression of key epigenetic regulators of histone acetylation and deacetylation in periodontal cells by quantitative PCR (**Fig. 1a, b**). BSA supplemented medium was used as a control. Compressive force was applied for six hours and non-stressed cells were used as controls. Yet, we could not detect any changes in genes encoding key histone acetylases *(TAF1, EP300, CREBBP)* and ubiquitously expressed histone deacetylases *(HDAC1-3),* neither due to fatty acid exposure nor to compressive stress.

**Figure 1.**
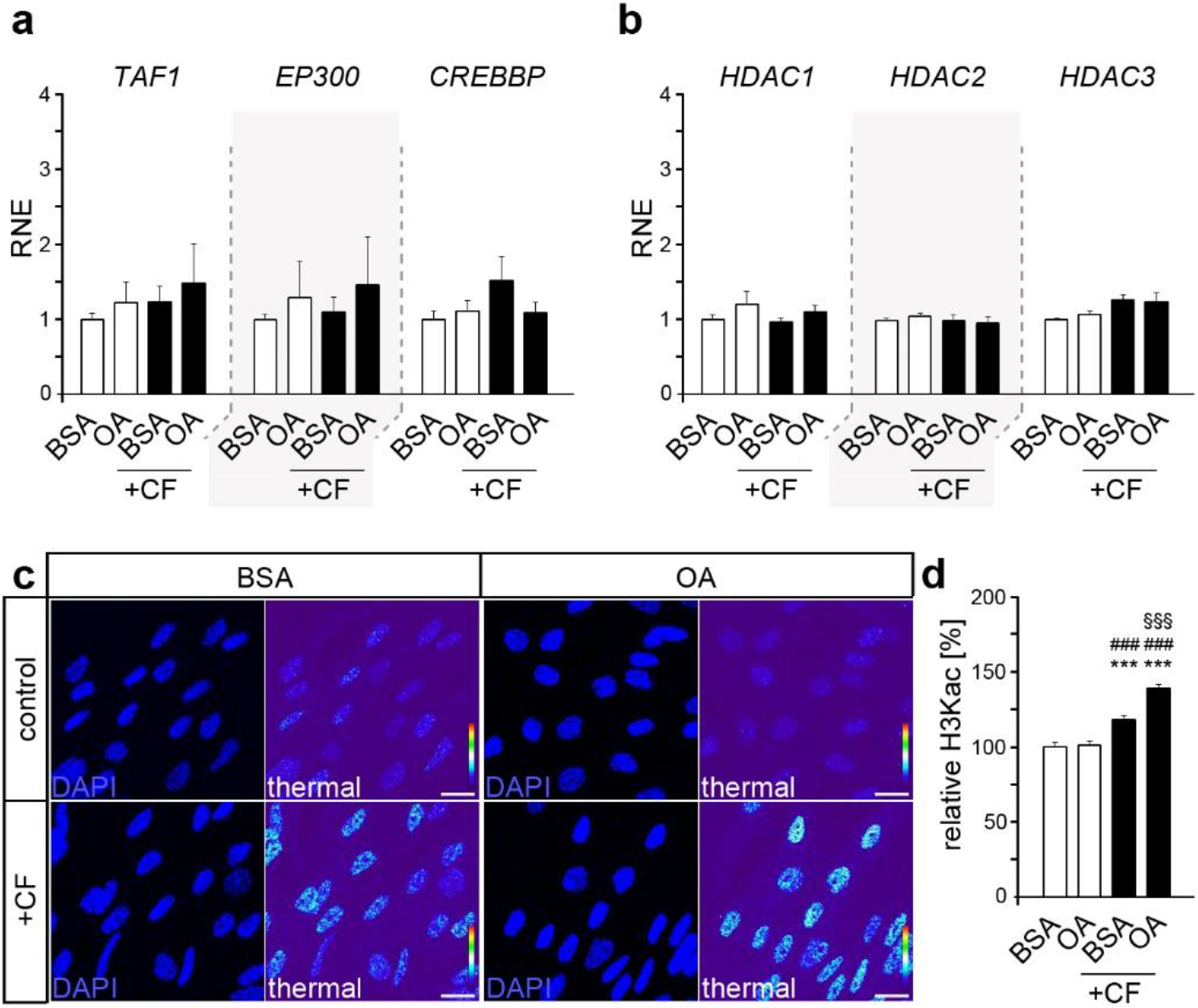
Compressive force increases global histone lysine acetylation further augmented by oleic acid exposure. (**a, b**) Quantitative expression analysis of genes encoding histone acetyl transferases *TAF1, EP300* and *CREBBP* (a) and encoding histone deacetylases *HDAC1, HDAC2* and *HDAC3* (b) in human periodontal ligament fibroblasts (HPdLF) exposed to oleic acid (OA) and stimulated with compressive force (CF) compared to BSA controls. (**c, d**) Analysis of global H3K9/14/18/23/27 acetylation (H3Kac) level showing representative microphotographs of HPdLF treated with OA and stimulated with CF in comparison to BSA controls (c). DAPI labels the cell nuclei and H3Kac staining is shown in thermal LUT (thermal), indicating intensity, which is analyzed in (d) in relation to BSA control. *** p < 0.001 in relation to BSA, ### p < 0.001 in relation to PA, §§§ p < 0.001 in relation to BSA + CF; One-Way ANOVA and post hoc test (Tukey). Scale bars: 10 μm in (c). RNE, relative normalized expression.

Semi-quantitative analysis of immunofluorescently labeled H3K9/14/18/23/27ac was performed to reveal differences in the formation of this epigenetic modification due to fatty acids as well as compressive force (**Fig. 1c, d**). While baseline levels remained comparable, force-related increase in H3K9/14/18/23/27 acetylation was higher in OA cultures (47.48% ± 5.35) than in BSA control (28.24% ± 5.07; p value = 0.0098, **). Thus, these data suggest that changes in the acetylation of histone 3-lysines may be involved in the OA-induced changes in the inflammatory response of HPdLF to compressive forces.

### Histone lysine acetylation promote IL10 expression in HPdLF

To evaluate the role of histone acetylation and deacetylation processes for the force-induced inflammatory response of HPdLF, we first inhibited histone deacetylation activity with romidepsin, which is a classic HDAC inhibitor with wide application in research and clinical settings [46].

To avoid possible toxic effects, the influence of different inhibitor concentrations within the usual range of application on the total cell number after one day of application was determined by MTT assay (**Fig. 2a**). At all concentrations used, no toxic effects were observed in one-day treatment. However, even at the lowest concentration of 10 nM, a drastic reduction in metabolic activity was observed after more than 2 days of cultivation.

**Figure 2.**
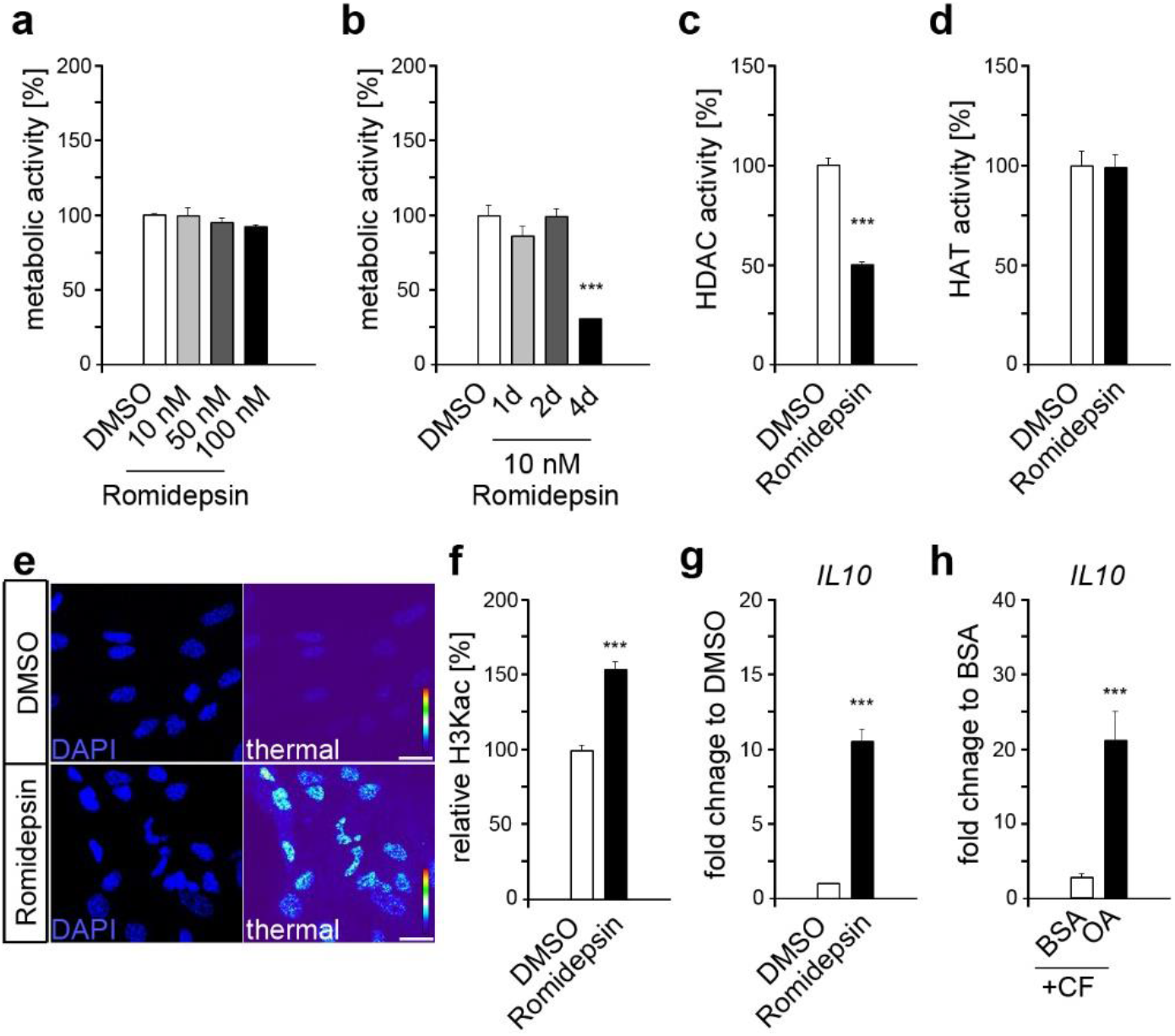
Histone lysine deacetylation revealed increased *IL10* expression comparable to that of compressed HPdLF exposed to oleic acid. (**a**) Analysis of the metabolic activity of human periodontal ligament fibroblasts (HPdLF) treated one day with different concentrations of romidepsin, a histone deacetylase (HDAC) inhibitor. DMSO was used as control. (**b**) Analysis of metabolic activity of HPdLF treated for 1d, 2d or 4d with 10 nM romidepsin compared to DMSO control. (**c, d**) Analysis of the activity of HDAC’s (c) and histone acetyltransferases (HAT; d) in HPdLF treated one day with 10 nM romidepsin in comparison to DMSO control. (**e, f**) Analysis of global H3K9/14/18/23/27 acetylation (H3Kac) level showing representative microphotographs of HPdLF treated one day with 10 nM romidepsin compared to DMSO control (e). DAPI labels the cell nuclei and H3Kac staining is shown in thermal LUT (thermal), indicating intensity, which is analyzed in (f) in relation to DMSO control. (**g, h**) Quantitative expression analysis of *IL10* in HPdLF treated one day with 10 nM romidepsin (g) as well as in HPdLF exposed to oleic acid (OA) and stimulated with compressive force (CF) in comparison to BSA control. Expression data in (f) were normalized to the non-forced BSA control (not shown). *** p < 0.001 in relation to DMSO in (b), (c), (f) and (g), and in relation to BSA in (h); One-Way ANOVA and post hoc test (Tukey). Scale bars: 10 μm in (e).

The efficacy of romidepsin was evaluated by HDAC inhibition assays, which revealed a 49.09% ± 3.45 inhibition even at 10 nM (**Fig. 2c**). For these reasons, we chose this concentration for the following experiments. One-day treatment with 10 nM romidepsin showed no direct effect on histone acetyltransferase activity (**Fig. 2d**), but resulted in increased H3K9/14/18/23/27ac levels (**Fig. 2e, f**). In this way, the force-dependent increase in H3K acetylation was simulated, but without a variety of further changes due to compressive stress. Quantitative expression analysis revealed a significant increase of *IL10* expression in HPdLF exposed to romidepsin (**Fig. 2g**). *IL10* transcription was also strongly enhanced in force-stressed HPdLF treated with OA compare to BSA control (**Fig. 2h**) suggesting it as potential candidate for the anti-inflammatory effect of OA, which expression can be modulated by histone acetylation.

### OA-related anti-inflammatory effects in force-stressed HPdLF depends on H3K acetylation of IL10 promoter regions

To evaluate whether the OA-induced increase in H3K acetylation has an anti-inflammatory effect, we reduced H3K acetylation by specific HAT inhibitors. For this purpose, we used anacardic acid with a broad specificity and C646 as a specific p300/CBP inhibitor [47,48]. Both compounds had a positive effect on HPdLF growth even at low concentrations (**Fig. 3a**), reliably decreased HAT activity (**Fig. 3b**), and reduced acetylation of H3K9/14/18/23/27 (**Fig. 3c**). Applying a THP1 adhesion assay, we analyzed the inflammatory response of HPdLF triggered by the specific cultivation conditions. Once sensing pro-inflammatory signals, these non-adherent monocytic cells can adhere to the appropriate site and differentiate [49]. Both inhibitors resulted in reduced numbers of adherent THP1 cells in the force-stressed BSA control (**Fig. 3d, e**) suggesting general anti-inflammatory properties. Nevertheless, further OA-related anti-inflammatory properties in compressed HPdLF appear to be prevented by both AA and C646, as THP1 cell adhesion was comparable to BSA control. Quantitative PCR revealed down-regulated *IL10* expression in force-stressed HPdLF treated with both AA and C646 (**Fig. 3f**). Together, our data implies that IL10 can be modulated by H3K acetylation and that p300/CBP has a major role in this regulation in HPdLF.

**Figure 3.**
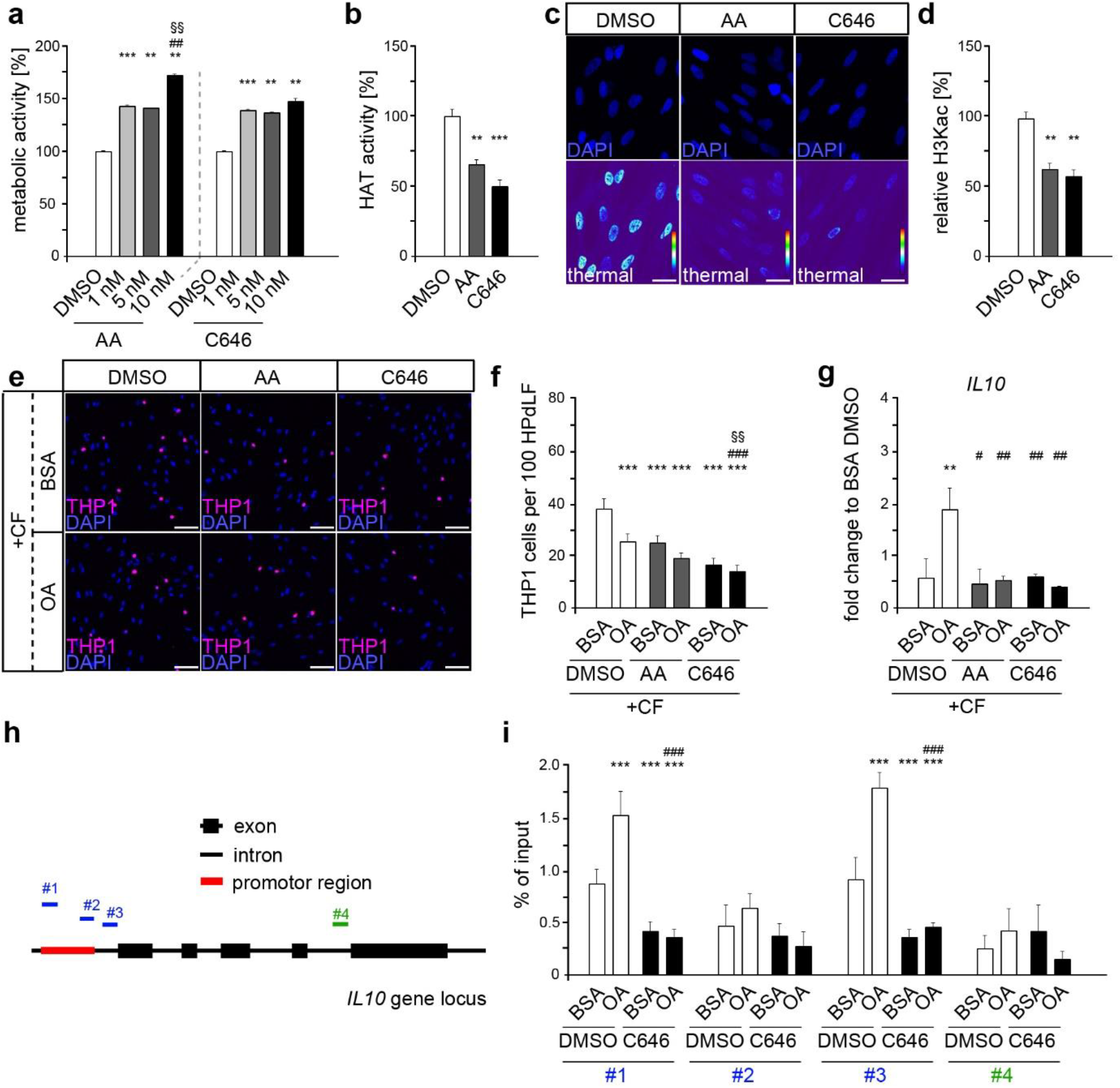
Oleic acid exerts its anti-inflammatory effect in force-stressed HPdLF via increased histone acetylation potentially on the *IL10* gene promoter. (**a**) Analysis of the metabolic activity of human periodontal ligament fibroblasts (HPdLF) treated one day with different concentrations of anacardic acid (AA) or C646, both histone acetyltransferase (HAT) inhibitors. DMSO was used as control. (**b**) Analysis of HAT activity in HPdLF treated one day with 10 nM AA or 1 nM C646 in relation to DMSO control. (**c, d**) Analysis of global H3K9/14/18/23/27 acetylation (H3Kac) level showing representative microphotographs of HPdLF treated one day with 10 nM AA or 1 nM C646 compared to DMSO control (c). DAPI labels the cell nuclei and H3Kac staining is shown in thermal LUT (thermal), indicating intensity, which is analyzed in (d) in relation to DMSO control. (**e, f**) Analysis showing microphotographs of fluorescently labeled, adherent THP1 monocytic cells (violet) on compressed HPdLF exposed to oleic acid (OA) after one-day treatment with 10 nM AA or 1 nM C646 in relation to BSA and DMSO controls (e) analyzed as relative number of THP1 cells per 10^2^ HPdLF in (f). Cell nuclei were labeled with DAPI. (**g**) Quantitative expression analysis of *IL10* in compressed HPdLF exposed to OA after one-day treatment with 10 nM AA or 1 nM C646 in comparison to BSA and DMSO controls. Expression data were normalized to the non-forced BSA DMSO control (not shown). (**h**) Locations of DNA primer amplified regions in the human *IL10* gene locus indicating promoter-associated pairs (blue, #1, #2 and #3) and a non-promoter-associated pair (green, #4). (**i**) Quantitative analysis of the association of specific *IL10* gene regions shown in (e) with acetylated H3K9/14/18/23/27 in compressed HPdLF exposed to OA and treated for one day with 1 nM C646 in comparison to the respective controls. Data were normalized to the sample input controls. ** p < 0.01, *** p < 0.001 in relation to DMSO in (a), (b) and (c) and in relation to BSA DMSO + CF in (f), (g) and (i); ## p < 0.01 in relation to 1 nM AA in (a); # p < 0.05, ### p < 0.001 in relation to OA DMSO + CF in (f), (g) and (i). §§ p < 0.01 in relation to 5 nM AA in (A) and in relation to BSA C646 + CF in (f); One-Way ANOVA and post hoc test (Tukey). Scale bars: 10 μm in (c) and (e).

To determine whether *IL10* gene could be a direct target of OA-modulated H3K acetylation, we performed CHIP-qPCR to assess its association to acetylated H3 lysines. To this end, we analyzed tree different positions in the *IL10* promoter region (#1-3, blue) as well as one nonpromoter region (#4, green) as control (**Fig. 3g**). We detected enhanced association of H3K9/14/18/23/27ac in the promoter region of positions #1 and #3 of the force-stressed OA cultures compared with the respective BSA control, whereas #2 and control #4 were not affected. Treatment with C646 significantly reduced this association, further supporting the important role of H3K acetylation in the anti-inflammatory effects of OA, particularly via regulation of *IL10* expression.

## Discussion

This study examined the potential effects of oleic acid on force-induced changes in histone lysine acetylation that attenuate the pro-inflammatory response of human periodontal ligament fibroblasts to mechanical stress. Compressive force increased the global level of H3K acetylation, and exposure to OA further enhanced this force-induced increase in global H3Kac. Moreover, *IL10* promoter-specific association with acetylated H3 lysines was also increased in compressed HPdLF exposed to OA. Thus, the anti-inflammatory effects of OA in compressed HPdLF appear to be modulated, at least in part, by p300/CBP-modulated H3K acetylation resulting in increased *IL10* expression.

The influence of obesity on orthodontic tooth movement has been investigated in only a few studies, partially leading to controversial results. In prospective controlled clinical trials, Jayachandran *et al.* [38] reported a decreased rate of OTM associated with increased BMI in adolescents, whereas Saloom *et al.* [40] demonstrated increased rates of OTM and changes in PdL tissue response in obese patients. These increased rates of OTM might enhance the risks for root resorption or even tooth loss. However, the cellular effects of obesity-induced changes in PdL functionality related to mechanically induced inflammation and tissue remodeling have shown that obesity-induced enhanced levels of specific fatty acids such as oleic acid have potential effects on PdL properties [33,42]. Although this fatty acid is elevated in obese patients [9,50], a diet rich in OA has been shown to positively influence disease progression [32]. This may be due to its ability to counterbalance the effects of saturated fatty acids including palmitic acid [11,20,24,27].

In addition to direct binding and post-translational modifications of transcription factors, fatty acids can regulate gene expression by altering epigenetic modification such as histone marks [51,52]. This has been specifically shown in glucose-starved C2C12 mouse myoblast cells, which showed up-regulated H3K23 acetylation when supplemented with 200 μM oleic acid [53]. In human PdL fibroblasts, we found increased acetylation of H3K9/14/18/23/27 only when they were additionally stimulated with mechanical stress but not without. In contrast to Jo *et al.* [53], who used medium with low glucose content, we continuously cultured in a medium with high glucose. Starvation also makes HPdLF more sensitive to additional stimuli [54]. However, we can only speculate on what impact additional prior starvation may have had on global H3K acetylation levels.

Nevertheless, independent of fatty acid exposure, compressive force alone also triggered an increase in H3K acetylation in HPdLF, supporting comparable results in bone marrow mesenchymal stem cells (MSC) [55]. In this study, perpendicular compressive forces on parallel-aligned cells triggered decreased HDAC activity leading to increased histone acetylation. This pathway, presumably mediated by the nuclear matrix protein lamin A/C [55], may be of interest because PdL cells also exhibit robust lamin A/C expression that is even increased in a force-dependent manner for lamin A [56]. In this aspect, in several cell types oleic acid treatment triggered the formation of lamin A/C-positive nuclear tubules [57], which are important for calcium homeostasis that can considerably affect histone acetylation [58]. Thus, changes in lamin A/C expression or function could be accountable for the increased H3K acetylation that we detected in compressed HPdLF treated with OA. Future studies could focus on the specific intracellular and intranuclear alterations of specific signal transduction pathways and transducers caused by simultaneous exposure to fatty acids and mechanical forces.

Although we did not measure HDAC activity in compressed OA-cultured HPdLF, the comparable *IL10* expression increase triggered by romidepsin-mediated HDAC inhibition suggests that it could be sufficient for an anti-inflammatory response to restrict the activity of those epigenetic erasers. This seems to be in accordance with HDAC inhibition studies in several cell types such as bone marrow–derived antigen-presenting cells [59], RAW264.7 macrophages [60] as well as KB31, C2C12, and 3T3-J2 cell lines [61]. However, a wide variety of HAT inhibitor studies suggests that IL10 is also directly regulated by HAT activity [62–66]. This was also evident here, as mechanically stressed HPdLF cultured in OA no longer exhibited elevated *IL10* levels after HAT inhibition with anacardic acid and p300/CBP-specific C646. Although both inhibitors have anti-inflammatory properties [67], which we also demonstrated, they have previously been shown to negatively regulate IL10 expression and secretion [63,65,66]. Our data now suggest that the anti-inflammatory effects of OA in force-loaded HPdLF are mediated by p300/CBP since its inhibition was sufficient to prevent OA-related reduced THP1 cell adhesion and enhanced lysine acetylation at the *IL10* promoter. This seems not unlikely in view of the fact that C646 competes with acetyl-CoA for the p300 Lys-CoA binding pocket [48]. Since alterations in acetyl-CoA concentrations could affect p300 histone acetylation activity [68,69], increased availability of acetyl-CoA by OA exposure should thereby no longer have an impact. The general and OA-specific reduction in histone acetylation at *IL10* promoter sites further supports this.

The results of our study are limited by several reasons. Because of the complex metabolic influences that individual fatty acids may have, *in vivo* it is not only the excess of one fatty acid that is decisive for hyperlipidemia-related health problems, but in particular the ratio to other fatty acids. This complicates *in vitro* studies such as ours. Additionally, we performed our analyses only on a certain concentration of OA and mechanical stimulation. However, studies show that this concentration is in the range also detected in the serum of obese patients [12–14,28,70] and achieved by increased intake through OA-rich diet [71]. After six hours of mechanical stimulation, the early phase of the inflammatory response can be seen. Moreover, examination of a single H3 lysine acetylation rather than an average of H3K9, K14, K18, K23, and K27 could provide further insight into the specific changes that OA might induce. In addition, further analysis of other anti-inflammatory cytokines could provide a more comprehensive view of the changes associated with OA, which was, however, beyond the scope of the study.

In summary, our data highlight an important influence of oleic acid on the acetylation profile of histone lysins in human PdL fibroblasts under mechanical compression. Thus, also in PdL tissue, epigenetic modifications and associated transcriptional changes are associated with metabolic shifts as seen in obese patients, which may influence tooth movement. Considering that epigenetic inhibitors already used in a variety of diseases, this might be a possibility to reduce obesity-related orthodontic impairments and the associated risks for root resorption and bone loss.

## Authors Contributions

LS performed experiments and analysis as well as wrote the original manuscript draft; KvB performed experiments; CJ revised the manuscript draft and contributed to project supervision; JS designed the project, acquired funding, supervised the project, performed experiments, figure illustration and revised the manuscript draft. All authors have read and agreed to the published version of the manuscript.

## Data availability statement

The datasets of this study are available from the corresponding author on reasonable request. The data are not publicly available due to very large size of microscopy images.

